# Topological Structures in the Space of Treatment-Naïve Patients With Chronic Lymphocytic Leukemia

**DOI:** 10.1101/2024.05.16.593927

**Authors:** Reginald L. McGee, Jake Reed, Caitlin E. Coombes, Carmen D. Herling, Michael J. Keating, Lynne V. Abruzzo, Kevin R. Coombes

## Abstract

Patients are complex and heterogeneous; clinical data sets are complicated by noise, missing data, and the presence of mixed-type data. Using such data sets requires understanding the high-dimensional “space of patients”, composed of all measurements that define all relevant phenotypes. The current state-of-the-art merely defines spatial groupings of patients using cluster analyses. Our goal is to apply topological data analysis (TDA), a new unsupervised technique, to obtain a more complete understanding of patient space. We applied TDA to a space of 266 previously untreated patients with Chronic Lymphocytic Leukemia (CLL), using the “daisy” metric to compute distances between clinical records. We found clear evidence for both loops and voids in the CLL data. To interpret these structures, we developed novel computational and graphical methods. The most persistent loop and the most persistent void can be explained using three dichotomized, prognostically important factors in CLL: *IGHV* somatic mutation status, beta-2 microglobulin, and Rai stage. In conclusion, patient space turns out to be richer and more complex than current models suggest. TDA could become a powerful tool in a researchers arsenal for interpreting high-dimensional data by providing novel insights into biological processes and improving our understanding of clinical and biological data sets.

**Simple Summary:** Clinical data sets incorporate continuous data like blood pressure or sodium levels, categorical data like cancer grade or stage, and binary data like sex or marital status. Measurements on an individual patient define a point in a high-dimensional space; data from many patients defines a “point cloud”. The “shape” of the point cloud influences experimental design by describing patient variability. Topological data analysis (TDA) is a mathematical technique for understanding the shape of point clouds by finding “holes” that correspond to combinations of patient characteristics that are never observed. TDA results are stratified by dimension. Zero-dimensional features define patient subtypes. One-dimensional features (“loops”) are analogs of the inside of a circle or a donut hole. Two-dimensional features (“voids”) are analogs of the inside of a balloon. Here, we apply TDA to a clinical data set of previously untreated patients with Chronic Lymphocytic Leukemia to find loops and voids.

## Introduction

Perhaps the greatest challenge in the analysis of clinical data is that patients are heterogeneous, with both diverse phenotypes and rare outlier presentations that affect disease outcome and treatment response. Clinical data, whether from electronic health records, clinical trials, or epidemiological studies, consists of large-scale, complex data sets. Analyses of clinical data are further complicated by noisy records, missing data, and mixed-type (both continuous and categorical) data [1–8]. To address these challenges, we view the full spectrum of data for any given disease as a high-dimensional “space of patients”. This patient space is composed of all measurements that define all relevant patient phenotypes. In a clinical study, the ideal design should sample patients across the patient space in order to encompass patient heterogeneity. For example, when a statistician designs a clinical trial and wants to estimate the variance from preliminary data, they need to have a reasonable idea of the number and distribution of distinct phenotypes.

One goal for an analyst is to uncover structures in the data that describe clinically relevant phenotypes. A variety of computational techniques are available to interrogate the shape of patient space and uncover meaningful patterns. These techniques include classical clustering methods that find structure in the form of distinct patient subtypes. High-dimensional structures can also be reduced to low-dimensional visualizations to aid interpretation by a variety of algorithms, including multidimensional scaling (MDS), t-stochastic neighbor embedding (t-SNE), and uniform manifold approximation and projection (UMAP) [9–11]. To move beyond the simple groupings captured by cluster analyses, here we investigate the applications of “topological data analysis” (TDA) to a space of patients with Chronic Lymphocytic Leukemia (CLL).

In the early 2000’s, mathematicians introduced an idea that is now called TDA [12–15], implemented using a notion called persistent homology [16,17]. The idea was to take the concepts of homology from the mathematical fields of algebraic topology and algebraic geometry and apply them to real-world data. In pure mathematics, homology provides a simple set of algebraic tools to discover “holes” in a topological space. The zero-dimensional homology counts the number of connected components, generating well-separated clusters. One-dimensional homology counts the number of “loops”, represented by the inside of a circle that encloses a hole in patient space. Two-dimensional homology counts the number of “voids”, which are represented by the inside of a hollow sphere. One should note that holes are inseparable from the boundary that encloses them. You can only detect holes in data if there are actual observed data points around them. The dimension of the hole is also defined by the dimension of the boundary. For instance, the inside of a sphere cannot be encapsulated by a circle; it requires a two-dimensional boundary.

While clusters are related to distinct phenotypes, loops and voids allow the separation of phenotypes based on more subtle differences between patient groups, with potential implications for feature selection and prognosis. Furthermore, the empty space within a loop or void suggests a state uninhabited by patients, which may carry clinical or biological meaning. In order to extract that meaning, it is necessary to understand how the expression of different clinical features varies over the boundary around the hole. For example, a single-cell study (using either mRNA-Sequencing or mass cytometry) of proliferating cells would be expected to detect a one-dimensional hole corresponding to the cell cycle. Cell-cycle related proteins or genes, such as Ki-67, cyclins, or cyclin-dependent kinases, would be expected to change expression around the loop, peaking in different phases of the cell cycle. In that example, it would be natural to think of the loop being parametrized by time as cells move though the cell cycle. Two-dimensional voids are harder to interpret in a similar way. Since there are two directions in which to move around the surface of a sphere, they can’t both represent time. One or both might represent the concentration of a critical molecule or metabolite. In the clinical setting, it remains unclear how to interpret the boundaries of loops or voids. What is clear, however, is that the presence of a hole suggests some mechanism that prevents the combination of features that should be present at the center of the hole from actually arising in a single patient.

It is *a priori* unclear to what extent one should expect loops or voids to arise in a clinical data set. Although zero-dimensional homology has been applied for cluster generation in a variety of contexts [18–22], to the best of our knowledge, this paper contains the first application of one- or two-dimensional homology to study patient space.

Novel computational methods have previously been leveraged to gain medical insights or generate hypotheses from the space of patients. In Ba-Dhfari’s University of Manchester thesis [23], they created a pair-wise semantic similarity score matrix from patient Read codes, followed by dimension reduction using principal component analysis and clustering with DBSCAN. Ba-Dhfari was able to recapitulate established disease associations between diabetes and circulatory system diseases, among others. Their novel computational method created useful visualizations of the space of patients, which rediscovered disease associations with a solid grounding in the literature. Another study employed a generative machine-learning model (specifically, a Hidden Markov Model) trained on laboratory blood test results and vital signs of 11,158 patients [24]. They found that a specific state space was strongly associated with inpatient and post-discharge mortality. The characteristics of this state closely resembled the description of Systemic Inflammatory Response Syndrome. In addition, a majority of the states (71%) had at least one strong association with an ICD-10 code (chi-squared p-value <0.001 after Bonferroni correction).

There is also a long history of viewing patient phenotypes and biological outcomes as lying atop a landscape or, more generally, a surface or manifold [25,26]. Mathematical models have been used for supervised explorations of such landscapes in order to address questions regarding the molecular mechanisms underlying homeostasis. When considering variation of key molecules the response of dopamine maps out a surface that can be used to understand phenotypic differences [27]. These studies highlight the need for new computational methods that are able to produce insightful visualizations of large clinical data sets for the purpose of exploratory analysis and hypothesis generation.

CLL is the most common adult form of leukemia in the United States, with approximately 20,000 new cases each year, and over 4000 deaths [28]. Although the disease can usually be successfully treated, relapse occurs eventually in a majority of cases [29]. Numerous prognostic factors are well established, including the somatic mutation status of the immunoglobulin heavy chain variable region genes (*IGHV*), the presence of specific chromosomal abnormalities, and serum levels of beta-2 microglobulin (B2M), among others [30]. There are also well established clinical staging systems (Rai or Binet). One might imagine that “holes” in a set of clinical data from CLL patients could arise from missing or rare combinations of some of these factors. For example, it would be unusual for a newly diagnosed CLL patient to have mutated *IGHV* genes (good prognosis) with del(17p) and high Rai stage (both bad prognosis).

In this manuscript, we test the hypothesis that TDA can detect and identify “loops” or “voids” in a clinical data set of CLL patients. The specific data set in question contains information on 266 newly diagnosed CLL patients seen at the M.D. Anderson Cancer Center between 1999 and 2013. The patients were recruited under an IRB-approved protocol to explore connections between known clinical factors, outcomes, and various omics technologies. We have previously published analyses of different subsets of this cohort [6,31–34].

## Methods

### Data

The data set consists of 30 measured or derived clinical variables on 266 treatment-naïve patients with CLL. The clinical data were collected at the time a patient first presented at the University of Texas M.D. Anderson Cancer Center (MDACC). Because MDACC is a tertiary care center, many patients are diagnosed elsewhere and later referred to MDACC. Thus, the starting point for any time-to-event analysis is taken to be “at presentation” rather than “at diagnosis”. At the time when these patients presented, combination chemoimmunotherapy (fludarabine, cyclophosphamide, and rituximab; FCR) was the standard first line treatment but was not considered curative. Most patients achieved remission but ultimately relapsed. Since many patients were asymptomatic at presentation, treatment was withheld until symptoms developed in order to maximize the interval between presentation and relapse. Thus, being able to predict who would need earlier or later treatment (time-to-treatment) was a useful tool for clinicians.

All data were collected in a CLL research database initiated by one of us (MJK). Much of the data was collected before the adoption of an electronic medical record system. Thus, one of us (CDH) reviewed the original patient records to verify data accuracy before it was used in our study. Data collection was authorized by an IRB-approved protocol, consistent with the principles of the Declaration of Helsinki.

### Statistics

All statistical analyses, including the computation of summary statistics, were performed in R version 4.4.0 (2024-04-24 ucrt) of the R Statistical Software Environment [35]. Time-to-event (overall survival; time to treatment) analyses were performed using Cox proportional hazards models evaluated by the score (log-rank) test as implemented in version 3.5.8 of the survival R package [36]. For continuous variables, such as age at diagnosis, the hazard ratio (HR) represents the change in the hazard for each one-point increase in the variable. For binary variables, HR is computed for the second factor-level relative to the first factor-level.

Distances between pairs of CLL patients were computed using the daisy function in version 2.1.6 of the cluster R package [37]. Previous simulation studies have shown that daisy is the best method for computing distances on sets of mixed-type data [8]. Here “mixed-type” means a combination of continuous, binary (either symmetric or asymmetric), and categorical (either nominal or ordinal) data. Most clinical data sets, including the one we study here, consist of mixed-type data. To visualize how well individual samples were clustered, we use the silhouette width [38]. The silhouette width is a score between −1 and 1 that compares the average distance of a sample to other members of its own cluster with the average distance to the nearest other cluster. Widths near 1 indicate that the sample is well classified; negative widths indicate that it is poorly classified. Multiple visualizations of the daisy distance matrix were created using version 1.1.4 of the Mercator R package [39]. We also used two binary distance metrics (Pearson correlation, and Sokal-Michener; see [40] for details) from the Mercator package to compute distances between clinical binary features for clustering purposes. Topological data analysis (TDA) was performed using version 1.9.1 of the TDA R package. Statistical significance and visualizations of higher dimensional structures found by TDA were assessed using an empirical Bayes method implemented in version 0.6.2 of the RPointCloud R package. The empirical Bayes method is modified for this context from one originally introduced by Efron and Tibshirani to study differential expression with microarrays [41].

To find clinical features that explain a loop in the data, we first located its centroid and converted all features to numerical vectors (for example, binary factors to {0,1}). For each feature, we averaged the expression of all patients located in sectors about the centroid with angle width of 20 degrees. We modeled these averages as a sinusoidal function (linear functions of sine and cosine) in order to identify the ones best explained by circular behavior. We also defined a goodness-of-fit statistic, *κ* = MSE*/σ*^2^, that compares the mean squared error (MSE) from the model to the total variance of the data.

## Results

### Clinical Characteristics

We began by computing summary statistics of the clinical variables (**Table 1**). For continuous variables, such as age at diagnosis, the table shows the mean and standard deviation (SD). For categorical or binary variables, the table shows the number of patients in each category. Because some data values are unavailable for some patients, the table includes both the number of missing entries (“NA’s”) and the total number *N* of patients available for use in univariate time-to-event analyses. Some continuous variables (beta-2 microglobulin (B2M); lactate dehydrogenase (LDH); and white blood count (WBC)) are presented on both their original scale and after log-transformation. Many continuous variables have also been dichotomized into “High” or “Low” categorical (“Cat”) values using the standard cutoffs preferred by clinicians who treat CLL patients. These cutoffs are:

- B2M *≤* 4 mg/L (Low); B2M *>* 4 mg/L (High)
- WBC *≤* 150 *×* 10^9^/L (Low); WBC *>* 150 *×* 10^9^/L (High)
- CD38 expression *<* 30% (Low), CD38 expression *≥* 30% (High)

**Table 1:**
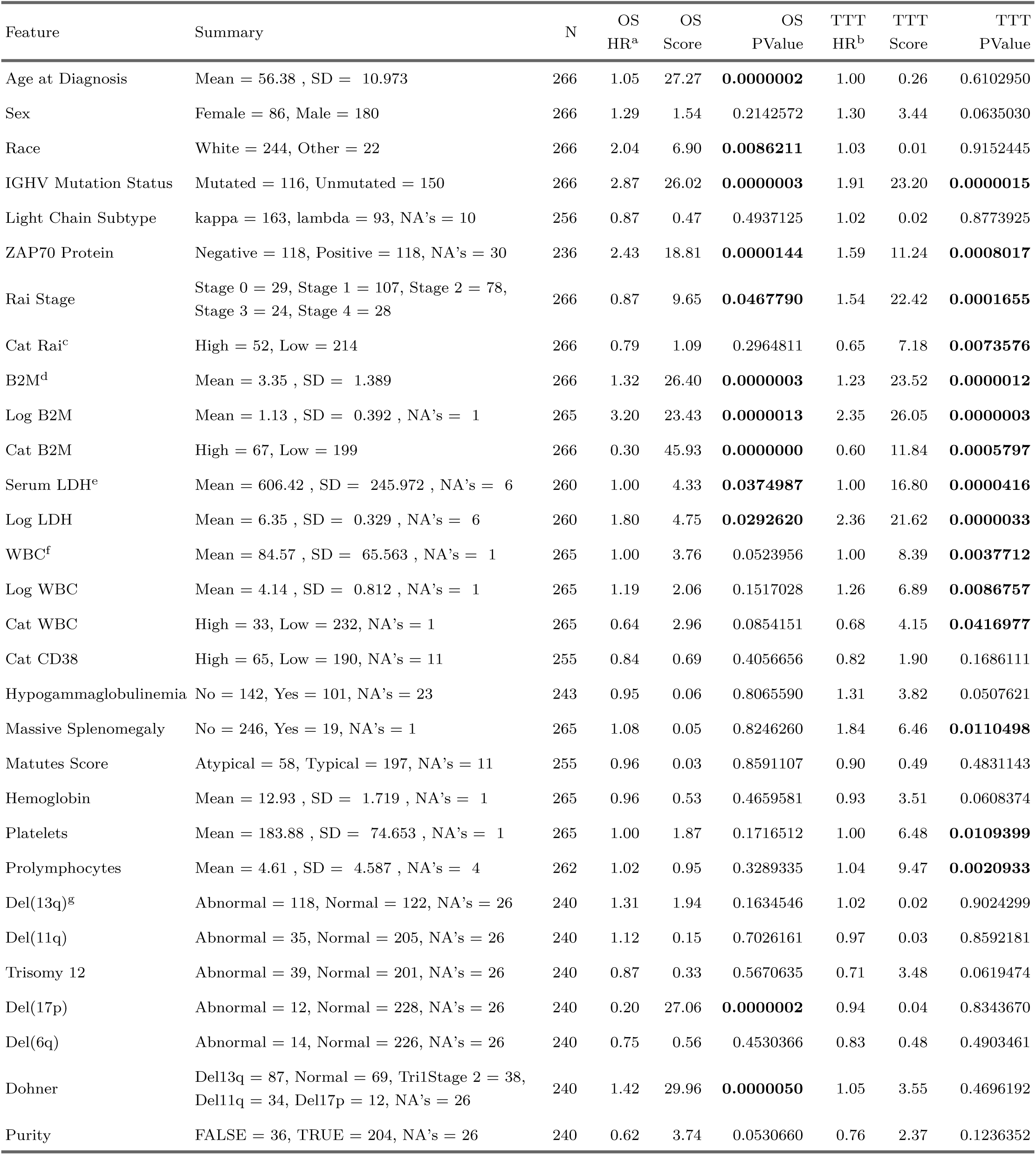
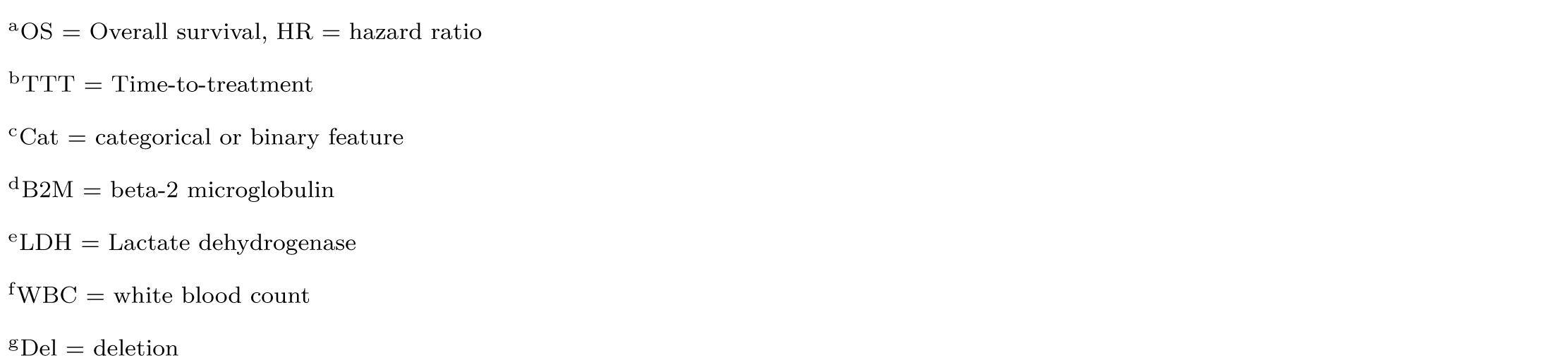
Clinical characteristics (p-values < 0.05 are shown in boldface).

### Time-to-event Analyses

**Table 1** displays the results of univariate Cox proportional hazards models to test the association between each clinical variable and two time-to-event (“survival”) outcomes. The first outcome tested was overall survival (OS), starting from the time each patient first presented at M.D. Anderson. For *IGHV* somatic mutation status, the hazard ratio HR = 2.87 indicates that *IGHV* -unmutated patients are almost three times as likely to die at any time after presentation as *IGHV* -mutated patients. The features most strongly associated with a difference in OS were B2M (as a binary variable, on the original continuous scale, or on the log-transformed scale), the presence of a 17p deletion (Del(17p)), age at diagnosis, *IGHV* somatic mutation status, the Döhner hierarchy of cytogenetic abnormalities, and a positive finding for the levels of ZAP70 protein.

When testing time-to-treatment (TTT), the best individual predictors included B2M, *IGHV* mutation status, and ZAP70 protein from the list of factors predictive of OS, along with lactate dehydrogenase (LDH) on the original or log scale, and Rai stage as either a five-level categorical variable or as a binary variable (CatRAI: Low = 0,1,2; High = 3,4).

All the findings for both time-to-event outcomes are consistent with previously published known prognostic factors in CLL, indicating that our cohort is typical of most cohorts of CLL patients at presentation.

### Daisy Distance

We used multiple methods to visualize the daisy distance between CLL patient samples. First, we used hierarchical clustering to arbitrarily define eight clusters (**Figure 1A**). Visualizations of the silhouette widths (**Figure 1B**) showed that both the “purple” and “black” clusters are difficult to separate from other clusters. We then plotted both linear (MDS; **Figure 1C**) and nonlinear (UMAP; **Figure 1D**) mappings of the daisy distance matrix into two dimensions. When using arbitrary distance metrics (like daisy), MDS creates a projection that does the best possible job of preserving pairwise distances. UMAP does not preserve distances but tries to preserve the “shape” of the data structures in high dimensions by focusing on local rather than global relationships. Both the MDS plot and the UMAP plot suggest the possibility of a similar “hole” in the data. The empty space inside this hole can be seen at the center of the UMAP plot, and at the top and slightly to the left in the MDS plot. The green and light blue clusters are on one side, and the other colors (especially the dark blue and magenta) are on the other side of the hole. These plots suggest the possibility of a “loop” around the hole that might be discoverable using TDA.

**Figure 1:**
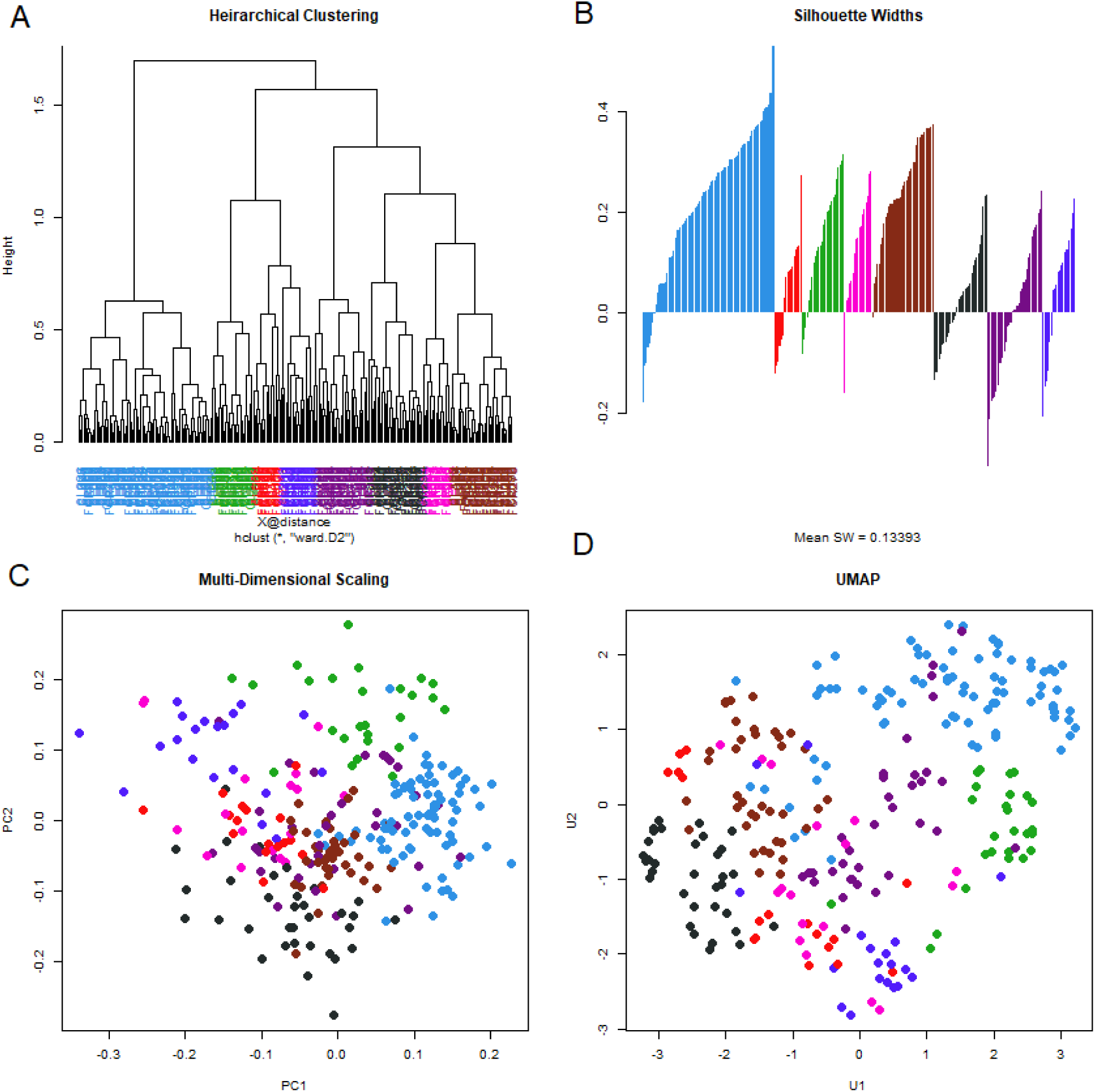
Visualizations of the clustered daisy distance matrix. (A) Hierarchical clustering was used to define eight clusters. (B) A silhouette width plot indicates which samples are well (positive score) or poorly (negative score) clustered. (C) Multi-dimensional scaling (MDS) linearly projects the data into two dimensions, preserving distances. (D) Uniform manifold approximation and projection (UMAP) creates a nonlinear mapping of the data into two dimensions, preserving shape but not distances.

### Topological Data Analysis

Next, we applied the TDA algorithm to the daisy distance matrix. The results of such an analysis are usually presented in a “persistence barcode” diagram, as shown in **Figure 2A**. In our case, the scale of the diagram is driven by the long black barcode at the bottom that corresponds to viewing the entire data set as a single connected point cloud. **Figure 2B** contains a “bean plot” or “violin plot” of the distributions of the lengths of barcodes (i.e., their persistence) by dimension. Lengths are shown on a log scale. In **Figures 2C** and **2D**, we zoom in on the dimension one (loops) and dimension two (voids) portions of the barcode. In each panel, the black arrow highlights the most persistent (longest) barcode in each set.

**Figure 2:**
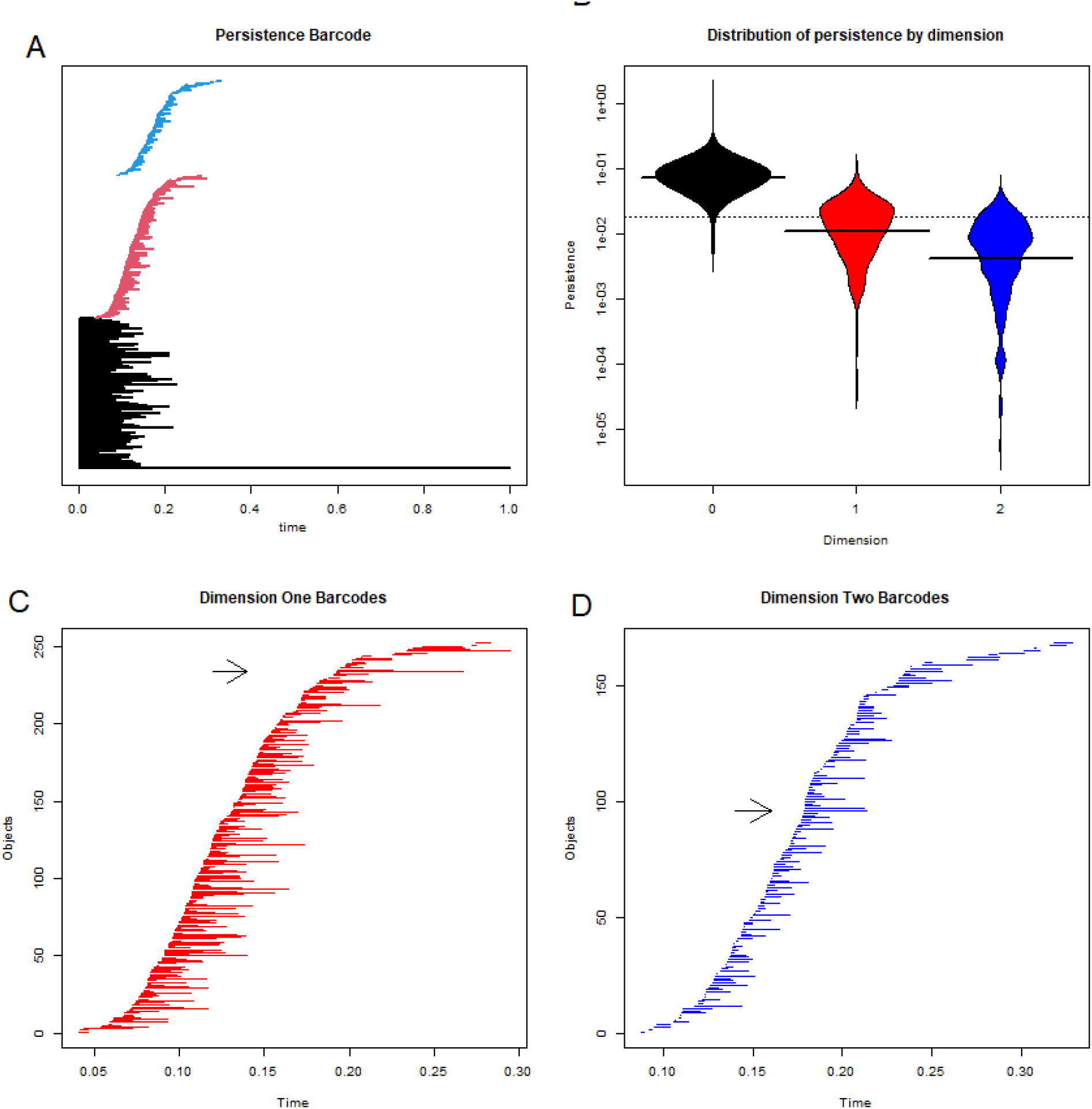
Persistence barcodes. (A) The standard plot of barcodes in all dimensions (black = 0, red = 1, blue = 2) is dominated by the long line that represents the point-cloud of the entire data set. (B) A beanplot of the distributions of (log) persistence by dimension. (C) Closeup of persistence barcodes in dimension 1. The black arrow highlights the longest barcode. (D) Closeup of persistence barcodes in dimension 2. The black arrow highlights the longest barcode.

### Statistical Significance of Loops and Voids

In order to test the statistical significance of the loops and voids found by TDA, we adapted an empirical Bayes approach first introduced into high-dimensional data analyses by Efron and Tibshirani during the early days of gene expression microarrays [41]. Visual inspection of the distributions of the durations of these features suggested that, for both loops and voids, they nearly followed an exponential distribution. Our unpublished simulations of unstructured data (from multivariate normal distributions) supported this idea. So, for the set of durations of observed structures in any dimension greater than zero, we robustly estimated the exponential parameter and modeled the data as a mixture of an exponential distribution (representing the null model) and an unknown distribution (representing the significant structures). **Figures 3A** and **3C** show these distributions for loops and voids, respectively. After choosing the most conservative possible prior parameters (which represent the probability that a random duration arises from the null model), we then computed the posterior probabilities that any duration arises from the alternative model (**Figures 3B** and **3D**). For the CLL data, this result implies that the most persistent loop and the most persistent void have a posterior probability of around 80%. (Other features may be significant at lower levels of posterior probability.)

**Figure 3:**
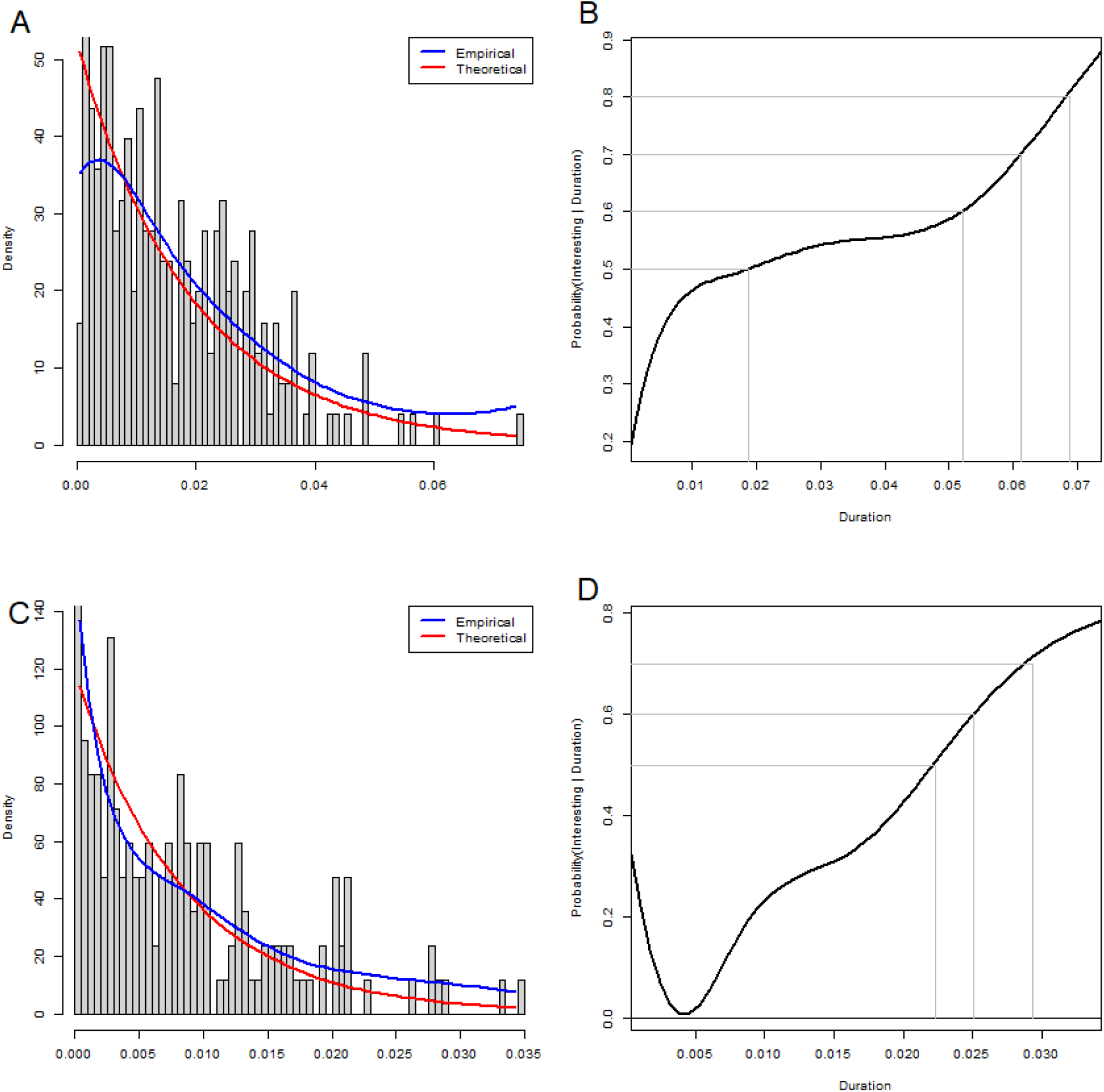
Empirical Bayes analysis of significance of loops and voids. (A) Histogram of loop durations (red = theoretical exponential distribution; blue = empirical distribution). (B) Posterior probability that a 1-cycle with this duration does not come from the exponential distribution. (C) Histogram of void durations. (B) Posterior probability that a 2-cycle with this duration does not come from the exponential distribution.

The most persistent loop and the most persistent void are found in the data set during overlapping time intervals. The loop exists during (0.193, 0.267), while the void exists during (0.179, 0.213).

### Interpreting Loops

We want to interpret the most persistent loop using information in the clinical data. We hypothesized that binary variables for which the two groups were widely separated would contribute to the formation of loops. (Continuous variables were dichotomized by splitting at the median to include them in this analysis.) So, for each binary variable, we computed the distance between the centroids of the two groups. The variables that defined the most widely separated groups were binary Rai stage (CatRAI, distance = 0.185), binary B2M (CatB2M, 0.171) and *IGHV* somatic mutation status (0.171). By comparison, all other factors had separations that were less than 0.14. We created multiple visualizations of the data, overlaying the support of the loop (blue curve) and the expression levels of binary clinical features (**Figure 4**). Black lines connect the centroids of the two levels. Each factor is concentrated at a different point around the loop, with a kind of “phase shift” from one feature to another.

**Figure 4:**
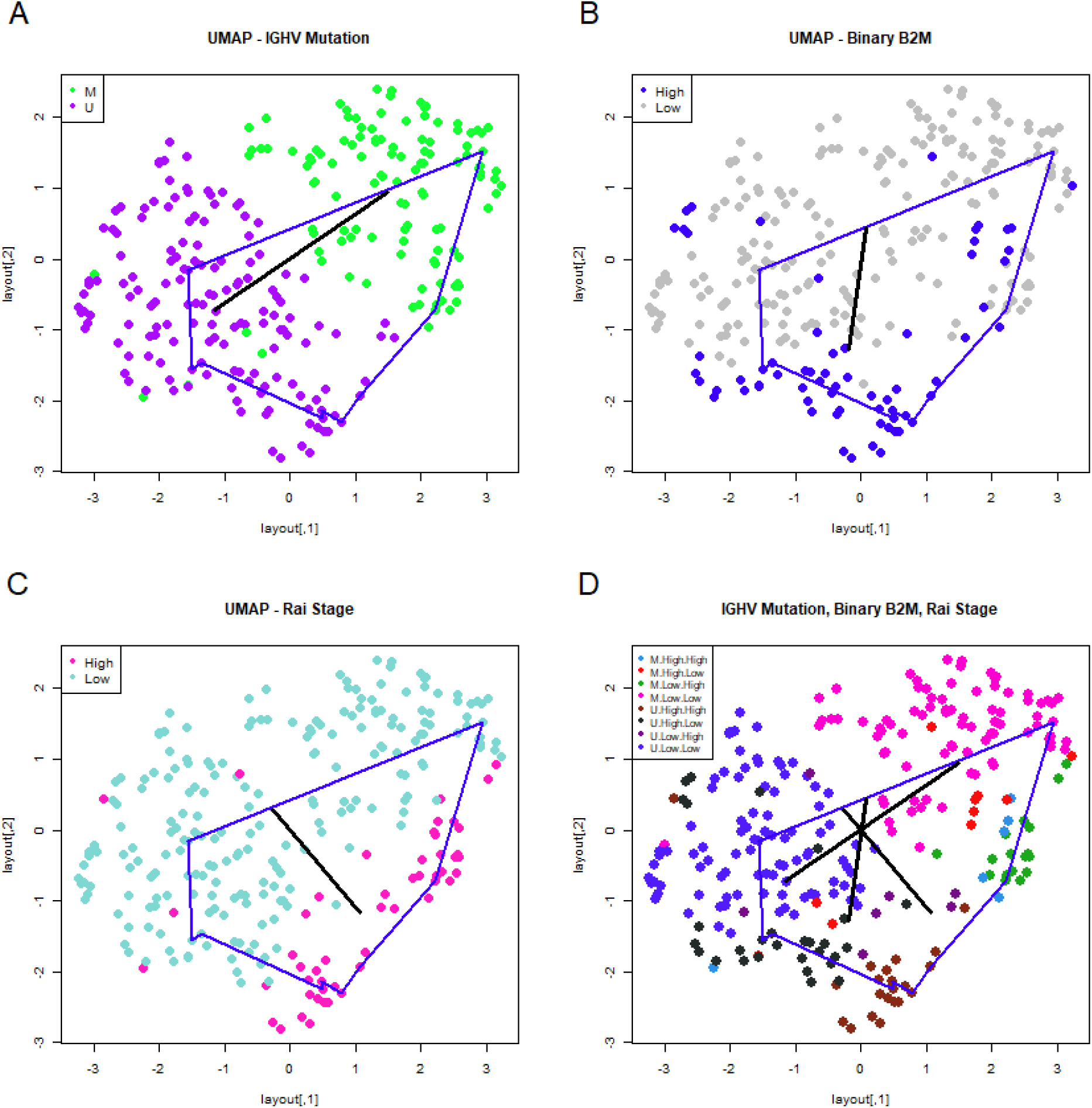
Interpreting the most persistent loop. In each panel, the blue curve connects the points identified by TDA as part of the most persistent loop. (A) UMAP plot with patients colored by *IGHV* mutation status. In this and the next two panels, the black line connects the centroids of the two subgroups of patients defined by this clinical feature. (B) UMAP plot colored by high or low levels of beta-2 microglobulin (B2M). (C) UMAP plot colored by binary Rai Stage. (D) UMAP plot colored by a combination of the three previous variables, showing all three centroid-connecting lines.

In **Supplemental Figure 1**, we show the expression patterns of four other binary features in the clinical data, each of which is associated to one of the three features shown in **Figure 4**. The “axes” connecting centroids of related features in **Figure 4** and **Supplemental Figure 1** (for example, *IGHV* mutation and ZAP70 expression) tend to point in similar directions. However, none of the features in **Supplemental Figure 1** are as clearly separated to opposite sides of the loop as those in **Figure 4**.

### Circos Plots

To find clinical features that best explain this loop in the data, we ranked the features by our goodness-of-fit statistic *κ*. Removing duplicates (in the form of different transformations of the same underlying data), the six most significant features were CatB2M, *IGHV* mutation status, CatRAI, the logarithm of LDH, the Döhner cytogenetic hierarchy, and hemoglobin levels. Plots of the fitted sinusoidal models are available in **Supplemental Figure 2**. In **Supplemental Figure 3**, we also present evidence that different features peak at different points around the loop. A circos plot of the combined expressions is shown in **Figure 5**. We can see overlaps between the poor prognosis regions associated with high levels of B2M, unmutated *IGHV* genes, high Rai stage, high levels of LDH, and “bad” cytogenetics according to the Döhner hierarchy.

**Figure 5:**
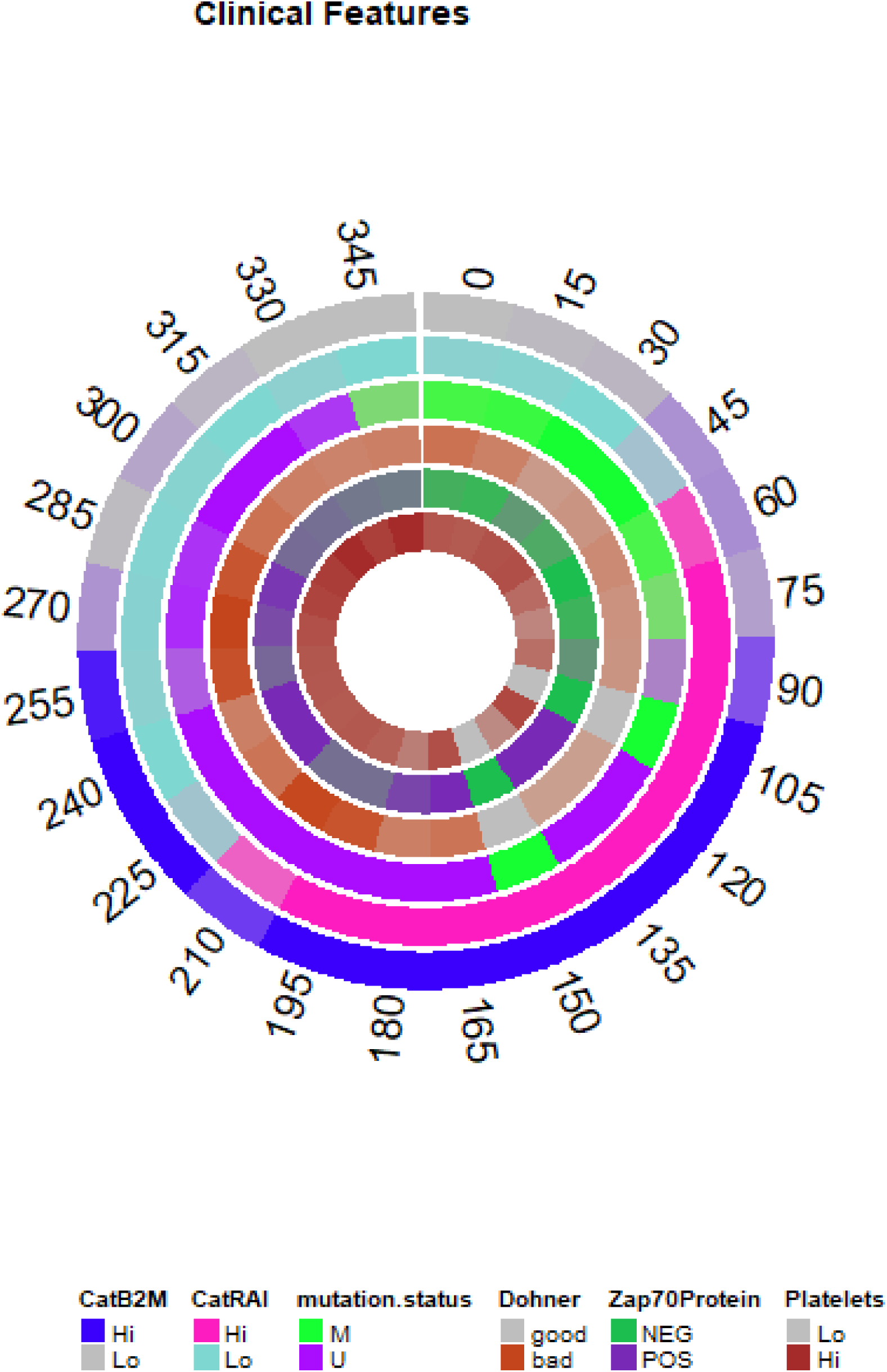
Circos plot of the mean expression of clinical features around the most persistent loop.

### Voids

The fact that the most persistent loop in the data can be described by three binary variables that point in three different directions suggests that the same variables may contribute to the presence of a void in this data set. In **Figure 6**, we present a three-dimensional view of the most persistent void in the data set. The three binary variables (*IGHV* mutation status, dichotomized B2M, and binary Rai stage) define a skewed coordinate system (shown in the figure in black). In **Supplemental Figure 4**, we also present “projection plots” for four of the clinical features, illustrating how expression changes on the surface of a unit sphere around the centroid of the void.

**Figure 6:**
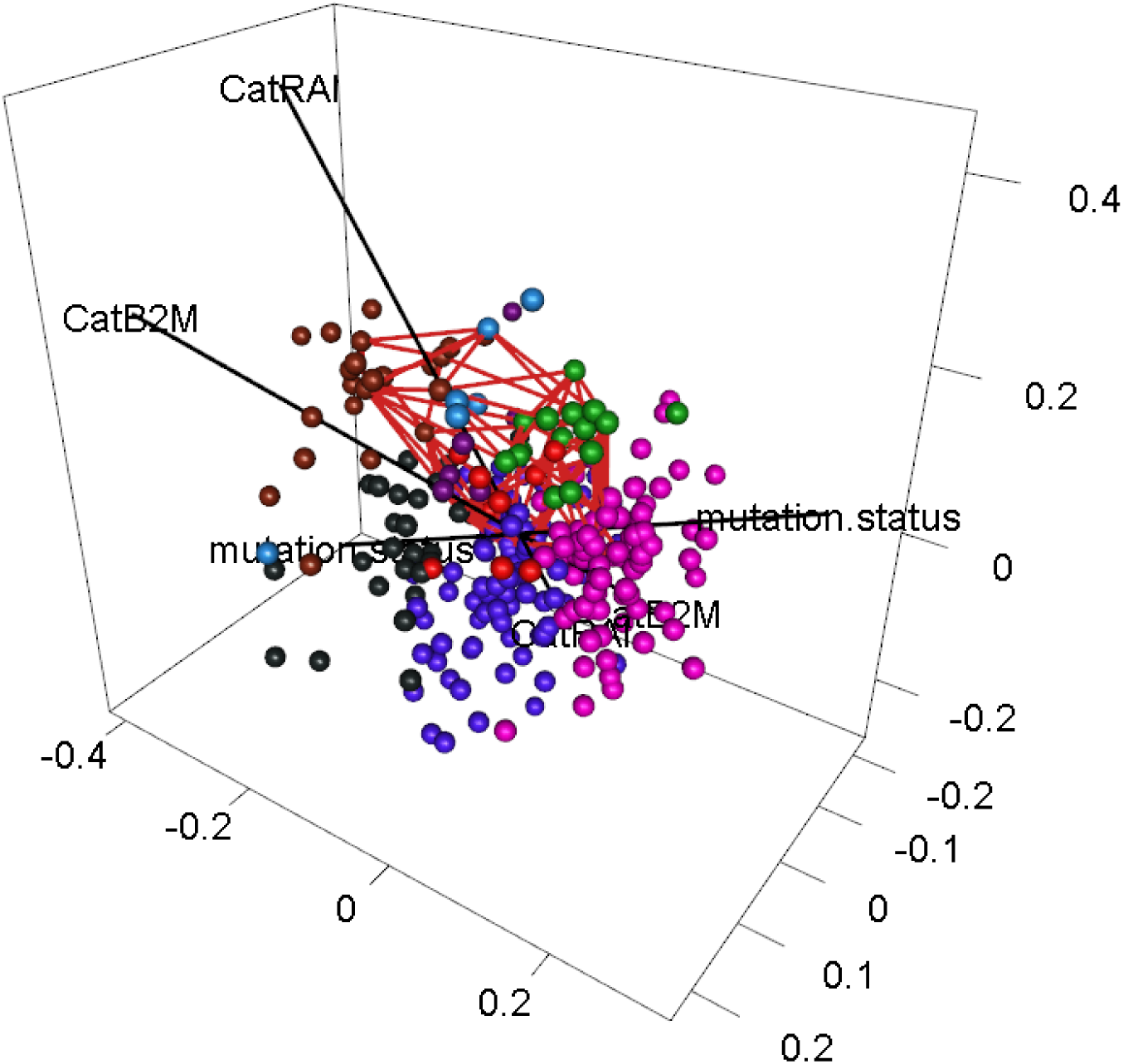
Three-dimensional plot showing the most persistent void (as a red skeleton) with three binary axes (mutation status, CatB2M, and CatRAI) in black. Patients are colored using the same scheme as Figure 3D.

### Dependence of Binary Features

The fact that we can interpret both the most persistent loop and the most persistent void as arising from three binary variables suggests the possibility that any collection of three such binary variables in clinical data should give rise to a void. However, we suspected that a stronger requirement may be necessary: the three variables should be independent. To test this idea, we performed chi-squared tests for each pair of binary features in the clinical data set. The negative log p-values from these chi-square tests are presented in a heatmap in **Figure 7**. The binary features were clustered as columns using Pearson correlation, and as rows using Sokal-Michener distance. We see three clusters of features that are found by both methods:

1. Dichotomous versions of serum beta-2 microglobulin (CatB2M) and Rai stage (CatRAI) are closely related (p = 3.257E-7).
2. Matutes score and trisomy 12 are closely related (p = 5.438E-8).
3. *IGHV* mutation status is closely related to ZAP70 protein levels (p = 4.456E-17).

**Figure 7:**
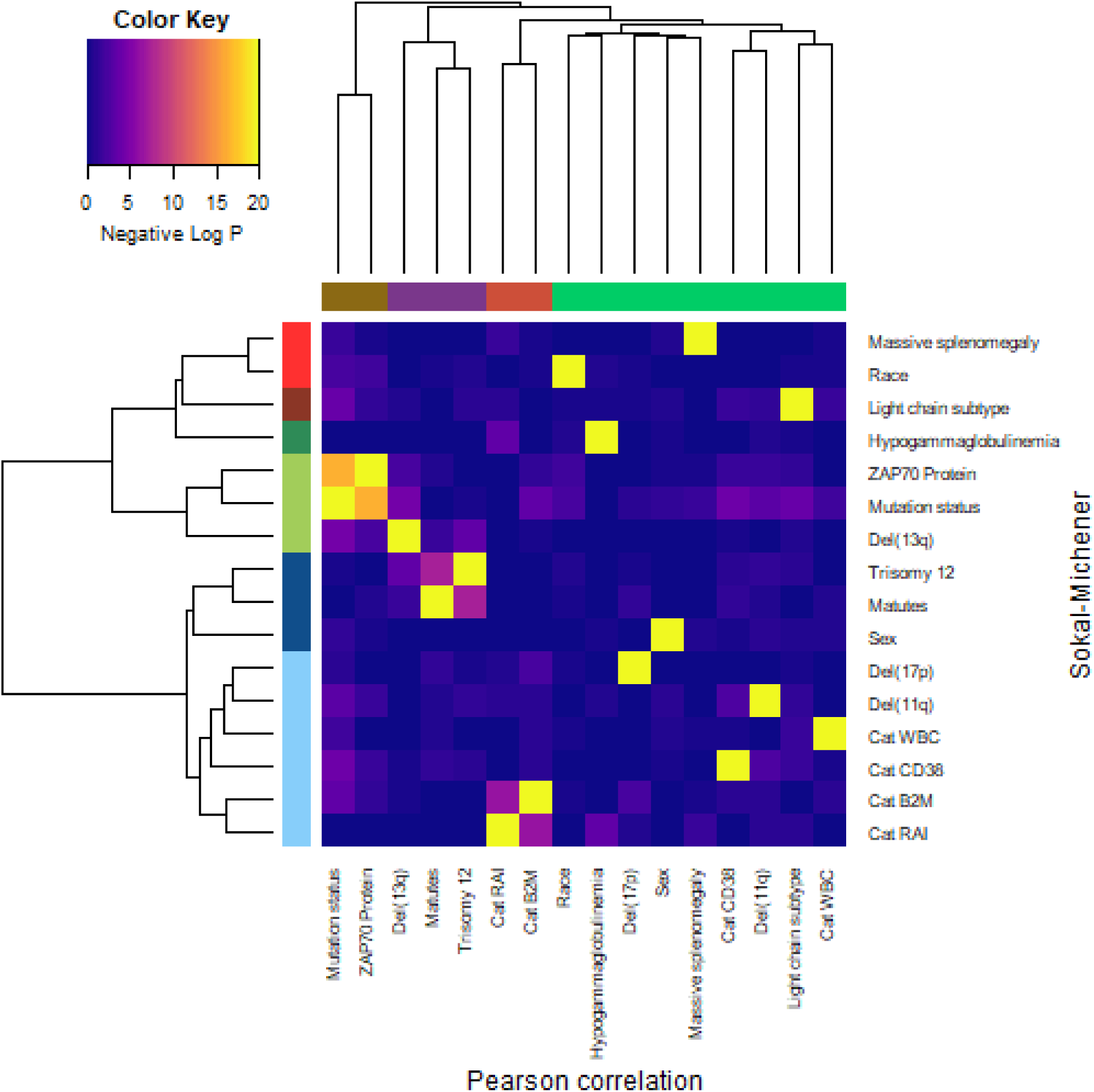
Heatmap and clustering of binary features in the CLL clinical data set.

We note that the del(13q) is somewhat related both to mutation status (p = 3.520E-5) and to trisomy 12 (p = 1.868E-4).

None of these findings are novel; their inclusion here illustrates the point that there are clear, strong dependency relationships between some of the binary features in the clinical data set. It is, however, somewhat surprising that the B2M and Rai stage variables appear to contribute (through linear independence) to the most persistent void in the data in spite of their strong statistical dependence.

## Discussion

In this paper, in order to better understand the underlying structure of “patient space”, we have applied a cutting-edge analysis tool, TDA, to a clinical data set containing 30 features assessed in 266 patients with CLL at their first presentation to the M.D. Anderson Cancer Center. The simplest model of patient space, derived from the idea that “all patients are similar” can be thought of as a single multivariate Gaussian distribution. The next simplest (and possibly most common) model, derived from the idea that “patients with this disease present with several distinct phenotypes”, can be thought of as a small collection of distinct multivariate Gaussian distributions, spatially well separated. Both simple models correspond topologically to a space with only zero-dimensional homology, which counts the number of distinct clusters or phenotypes.

Our application of TDA to the CLL data set shows that the true structure is richer and more complicated than those simple models. We have presented clear evidence for the presence of holes in the data, revealed by both loops and voids that can be described in terms of binary clinical variables. While it is true that each combination of binary values (for example, high Rai stage, high B2M, and unmutated *IGHV* genes) defines a specific phenotype, their arrangement in space (for example, centered at the vertices of a unit cube) gives rise to more interesting topology. One can imagine that the effect of other clinical variables in the higher dimensional space will “diffuse” the patient data near these vertices, thus occupying a larger portion of the surface of a cube that is, itself, topologically equivalent to a sphere. Clinically, this “diffusion” corresponds to the statement that the other clinical variables modulate the behavior of the main variables. In spite of our initial idea that the absence of patients with rare combinations of these variables might give rise to holes in the data, it instead appears that the presence of these rare cases helps sketch out the cubical/spherical outline of the holes.

In CLL, the somatic mutation status of the *IGHV* genes is known to be one of the most important prognostic features [42]. Mutation status is also associated with other features that have been proposed at various times as replacements or surrogates, including ZAP70 protein [43], the CD38 cell-surface marker [44], and immunoglobulin light chain usage [45]. Each of these factors is differentially present around the loop (and around the void) but define nearly the same axis of separation as *IGHV* mutation status.

Serum beta-2 microglobulin is another known prognostic factor [46], which often survives in multivariate linear models of time-to-event outcomes (like overall survival or time-to-treatment) as a predictor that is independent of *IGHV* mutation status. In our data, the dichotomized B2M levels define a second axis of separation. However, B2M is often associated with Rai stage, with low levels of B2M accompanying early stages (0, 1, or 2), and higher levels of B2M accompanying more advanced stage (3 or 4). Surprisingly, we found that these two statistically dependent binary variables produce linearly independent axes of separation, thus providing interpretable axes.

Another strong prognostic marker in CLL is defined by the Döhner hierarchy of cytogenetic abnormalities [47]. This hierarchy is defined by the presence of the first of the four most common abnormalities, ordered from worst to best prognosis: del(17p), del(11q), trisomy 12, or del(13q). Perhaps surprisingly, these abnormalities have a weaker association with our most persistent loop and void. Deletion 13q is associated with mutated *IGHV* genes, and deletion 11q is associated with unmutated *IGHV* genes, so both are indirectly present in the model through the inclusion of mutation status. However, deletion 17p did not show up, even though the clustered heatmap in **Figure 7** suggests that it is independent of everything else in the clinical data set. One possible explanation is that these abnormalities are rare in treatment-naïve patients. It is possible that del(17p) would show up as a “hole” in the data in higher dimensions, by contributing to a “hypersphere” in an ambient space of dimension 4 or more. We were unable to assess this possibility because of a fundamental computational limitation of current implementations of topological data analysis. The computational complexity of the computations grows combinatorially based on the number of points (i.e., patients) *M* in the data set and the dimension *D* of the hole being sought. To find loops, you only need to consider two points at a time to decide if they need to be connected by an edge. To find voids, you must consider all triples of points to decide if they should define a triangular “face”. To find higher dimensional holes (hypervoids?), you have to assess all combinations of four or more points at a time. Mathematically, the number of ways to select *D* points at a time out of *M* (known as *M* -choose-*D* and, for now, written as the function choose(M, D)) can be computed using factorials, and grows quite rapidly. Specifically, we see that choose(266, 2) = 35,245, and choose(266, 3) = 3,101,560, and choose(266, 4) = 203,927,570. We were unable to perform TDA when trying to assess four points at a time with a desktop computer that had 32GB of RAM.

The Matutes score was introduced originally as a tool to diagnostically distinguish CLL from other B-cell abnormalities by combining five surface markers measured by flow cytometry [48]. “Typical” CLL cases express all or most of the five markers; “atypical” cases express few or none. It was eventually realized that atypical cases are often associated with the presence of a trisomy 12 cytogenetic abnormality, which we also found in our data. In our data, the trisomy 12 patients are clearly not separated around the loop (**Supplemental Figure 1C**). The axis arising from Matutes score (not shown) is similar. It is possible that variables like Matutes score, Trisomy 12, and del(17p) contribute to other significant loops or voids in the data. Because of limitations of journal space, we have not made a deeper dive into the persistence barcodes to understand other potential loops and voids.

Readers who have paid close attention to this point may have noticed that the most persistent loop does not enclose the open area that we pointed out in the text initially describing **Figure 1C** and **1D**. There are such loops (examples are shown in **Supplemental Figure 5**), but their persistence/duration is significantly shorter than that of the loop we interpreted. Some readers may have also noticed that the inside of the most persistent loop in its 2D visualization is not empty, but encloses many data points (patients). These apparent oddities result because the TDA computations are performed in the full 30-dimensional space where the daisy distance was computed using the entire clinical data set. None of the 2D visualizations can fully preserve the distances or relative positions of all the points in 30 dimensions. If you imagine a sphere in 3D being projected into 2D, points near the north and south poles are among the ones that are furthest apart, yet their projections are among those closest together.

Even though multi-dimensional scaling is known to be optimal for preserving pairwise distances, one should keep in mind that this property refers to the *average* distances. The average is often achieved by drastically violating the property for some pairs of points. Both t-SNE and UMAP can distort the data in various ways. They are designed to preserve (most, but not all) local distances, and they provide no control over distances between pairs of points that are not close together in the original high-dimensional space. They can destroy topological structures by flattening them out, or by “tearing” the space in order to fit them into two dimensions. They can also create artifacts by arbitrarily expanding small gaps in the data into larger holes. One should keep these limitations in mind when evaluating other applications of dimension reduction already present in the biomedical informatics literature.

For the evaluation and interpretation of loops and voids, it may theoretically be possible to design individualized dimension reductions to better preserve the size and shape of an individual feature. We suspect that no single method will simultaneously work well for all loops and voids. Nevertheless, using TDA to explore patient space could offer benefits to generating virtual patient cohorts, which have recently proven to be a powerful analysis tool that can follow mathematical modeling of a biological process [49]. The detection of holes in patient space can give insight into sparse regions of patient space where certain feature combinations rarely or never occur. Reducing to only biologically observable combinations of features could help with efficiency in generating the virtual cohort and a better understanding of the variability and structure of the space could increase the utility of such analyses.

Although we were able to interpret one loop and one void using the binary features in the data, there is no reason to believe that this structure is the only way that such phenomena can arise. We see no reason why continuous data cannot give rise to similar structures. In fact, we are fairly confident that continuous single-cell measurements of the appropriate protein levels on proliferating cells should lead to a detectable loop corresponding to the cell cycle. It is even possible that the continuous versions of some of the markers in our data (such as the continuous measurements of serum B2M) helped make the topological structures detectable. Further research, with a wider variety of clinical data sets, still needs to be done.

## Conclusions

Topological data analysis is a powerful new tool to understand the structure of patient spaces defined by clinical data sets. We were able to detect loops and voids even in a relatively small, homogeneous clinical data set of CLL patients. Larger and less homogeneous data sets seem likely to include even more loops and voids, along with possibly higher dimensional holes. These topological structures take us far beyond the simpler common analyses that only detect phenotypes as clusters of related data points. The arrangements of clinical data in high-dimensions reflect the presence of holes of varying dimensions, each of which is likely to provide insight into important clinical and biological characteristics and their interactions.

To make these methods more accessible to researchers and informaticians, we have presented new tools (as part of the RPointCloud R package) to explore these structures. These include (i) empirical Bayes models to evaluate the statistical significance of loops and voids, (ii) a goodness-of-fit statistic to identify clinical features that contribute to the presence of loops through sinusoidal expression patterns, and (iii) multiple graphical tools to display loops, void, and features in two and three dimensions. We expect the combination of TDA and the RPointCloud package of tools will be of value to researchers trying to understand why and how these holes develop in the data that gives rise to clinically defined patient spaces.

## Supporting information

Suppplemental Figure 1

Suppplemental Figure 2

Suppplemental Figure 3

Suppplemental Figure 4

Suppplemental Figure 5

## Supplementary Materials

**Supplemental Figure 1:** Expression of binary features around the most persistent loop. In each panel, the blue curve connects the points identified by TDA as part of the most persistent loop, and the black line connects the centroids of the two subgroups of patients defined by this clinical feature. (A) UMAP plot with patients colored by Zap70 protein expression. (B) UMAP plot colored by high (3, 4) or low (0,1, 2) levels of Rai Stage. (C) UMAP plot colored by presence or absence of a trisomy 12. (D) UMAP plot colored by high or low levels of CD38.

**Supplemental Figure 2:** Goodness of fit modeling mean expression of clinical features around the most persistent loop in the CLL data. Observed data is shown as solid circles; the sinusoidal model as a dotted curve with open circles.

**Supplemental Figure 3:** Heatmap image of the fitted models of mean expression of clinical features around the most persistent loop in the CLL data. Data for each feature has been standardized so they are displayed on comparable scales.

**Supplemental Figure 4:** Projection plots of clinical feature expression around the surface of the most persistent void. To construct these plots, we first projected every data point (patient) onto the surface of a unit sphere around the centroid of the void. We then used the equivalent of a Mercator map projection to display the surface of the sphere in two dimensions, in terms of spherical coordinates of longitude and latitude. For each of four features (CatB2M, *IGHV* mutation status, CatRAI, and the Döhner cytogenetic hierarchy), the left-hand panel contains a scatter plot of the patients, colored by expression. The right-hand panel contains an image of the predicted expression from a two-dimensional smooth loess model of the observed values as a function of latitude and longitude.

**Supplemental Figure 5:** Other loops. The first three panels illustrate three loops in the CLL data that overlap the “obvious” empty area in the 2D UMAP visualizations. All three have considerably shorter duration that the most persistent loop (with duration > 0.07). The final panel is a smoothed scatter plot comparing the daisy distance in 30 dimensions to the Euclidean distance between points in the UMAP 2D plot.

## Author Contributions

Conceptualization, L.V.A. and K.R.C.; Data curation, C.D.H., M.J.K., and C.E.C.; Formal analysis, R.L.M., J.R. and K.R.C.; Methodology, R.L.M., J.R. and K.R.C.; Supervision, K.R.C. and L.V.A.; Writing—original draft, R.L.M. and K.R.C.; Writing—review & editing, L.V.A. All authors have read and agreed to the published version of the manuscript.

## Institutional Review Board and Informed Consent

This study uses deidentified data that were previously published. Originally, peripheral blood samples and clinical records were obtained from 266 treatment-naïve CLL patients after obtaining informed consent at the University of Texas MD Anderson Cancer Center. The studies were approved by the Institutional Review Board and conducted according to the principles expressed in the Declaration of Helsinki.

## Availability

The RPointCloud R package used to visualize and to test the significance of loops and voids can be obtained from the Comprehensive R Archive Network (https://CRAN.R-project.org/package=RPointCloud). The deidentified clinical data are included in the package as a binary R data file called CLL. The latest version of the package can always be obtained from R-Forge (https://r-forge.r-project.org/R/?group_id=2244). Full code to reproduce the analysis described in this manuscript is available in a Git repository (https://gitlab.com/krcoombes/cll-tda).

## Acknowledgments

None.

## References

[1] M. Juhola, J. Laurikkala, On distance computation in space of mixed-type variables in medical data mining., Studies in Health Technology and Informatics 90 (2002) 425–430.

[2] C.-C. Hsu, S.-H. Lin, Visualized analysis of mixed numeric and categorical data via extended self-organizing map., IEEE Transactions on Neural Networks and Learning Systems 23 (2012) 72–86. 10.1109/TNNLS.2011.2178323.

[3] M. Hummel, D. Edelmann, A. Kopp-Schneider, Clustering of samples and variables with mixed-type data, PloS One 12 (2017) e0188274.

[4] S. Zhang, J. Cao, C. Ahn, Sample size calculation for before-after experiments with partially overlapping cohorts., Contemporary Clinical Trials 64 (2018) 274–280. 10.1016/j.cct.2015.09.015.

[5] V. Cabeli, L. Verny, N. Sella, G. Uguzzoni, M. Verny, H. Isambert, Learning clinical networks from medical records based on information estimates in mixed-type data., PLoS Computational Biology 16 (2020) e1007866. 10.1371/journal.pcbi.1007866.

[6] C.E. Coombes, Z.B. Abrams, S. Li, L.V. Abruzzo, K.R. Coombes, Unsupervised machine learning and prognostic factors of survival in chronic lymphocytic leukemia, Journal of the American Medical Informatics Association: JAMIA 27 (2020) 1019–1027. 10.1093/jamia/ocaa060.

[7] S. Faisal, G. Tutz, Imputation methods for high-dimensional mixed-type datasets by nearest neighbors., Computers in Biology and Medicine 135 (2021) 104577. 10.1016/j.compbiomed.2021.104577.

[8] C.E. Coombes, X. Liu, Z.B. Abrams, K.R. Coombes, G. Brock, Simulation-derived best practices for clustering clinical data., Journal of Biomedical Informatics 118 (2021) 103788. 10.1016/j.jbi.2021.103788.

[9] I. Borg, P. Groenen, Modern multidimensional scaling: Theory and applications, Journal of Educational Measurement 40 (2003) 277–280.

[10] L. van der Maaten, G.E. Hinton, Visualizing high-dimensional data using t-SNE, J Machine Learning Rsch 9 (2008) 2579–2605.

[11] L. McInnes, J. Healy, N. Saul, L. Großberger, UMAP: Uniform Manifold Approximation and Projection, Journal of Open Source Software 3 (2018) 861. 10.21105/joss.00861.

[12] H. Edelsbrunner, D. Letscher, A. Zomorodian, Topological persistence and simplification, Discrete & Computational Geometry 28 (2002).

[13] G. Carlsson, A. Zomorodian, A. Collins, L.J. Guibas, Persistence Barcodes for Shapes, International Journal of Shape Modeling 11 (2005) 149–187.

[14] R. Ghrist, Barcodes: The persistent topology of data, Bulletin of the American Mathematical Society 45 (2008) 61–75. 10.1090/S0273-0979-07-01191-3.

[15] G. Carlsson, Topology and Data, Bull AMS 46 (2009) 255–308.

[16] A. Zomorodian, G. Carlsson, Computing Persistent Homology, Discrete & Computational Geometry 33 (2005) 249–274. 10.1007/s00454-004-1146-y.

[17] F. Iuricich, Persistence Cycles for Visual Exploration of Persistent Homology., IEEE Transactions on Visualization and Computer Graphics 28 (2022) 4966–4979. 10.1109/TVCG.2021.3110663.

[18] J. Bigler, M. Boedigheimer, J.P.R. Schofield, P.J. Skipp, J. Corfield, A. Rowe, A.R. Sousa, M. Timour, L. Twehues, X. Hu, G. Roberts, A.A. Welcher, W. Yu, D. Lefaudeux, B.D. Meulder, C. Auffray, K.F. Chung, I.M. Adcock, P.J. Sterk, R. Djukanović, A Severe Asthma Disease Signature from Gene Expression Profiling of Peripheral Blood from U-BIOPRED Cohorts., American Journal of Respiratory and Critical Care Medicine 195 (2017) 1311–1320. 10.1164/rccm.201604-0866OC.

[19] J. Brandsma, V.M. Goss, X. Yang, P.S. Bakke, M. Caruso, P. Chanez, S.-E. Dahlén, S.J. Fowler, I. Horvath, N. Krug, P. Montuschi, M. Sanak, T. Sandström, D.E. Shaw, K.F. Chung, F. Singer, L.J. Fleming, A.R. Sousa, I. Pandis, A.T. Bansal, P.J. Sterk, R. Djukanović, A.D. Postle, Lipid phenotyping of lung epithelial lining fluid in healthy human volunteers., Metabolomics : Official Journal of the Metabolomic Society 14 (2018) 123. 10.1007/s11306-018-1412-2.

[20] J.L. Bruno, D. Romano, P. Mazaika, A.A. Lightbody, H.C. Hazlett, J. Piven, A.L. Reiss, Longitudinal identification of clinically distinct neurophenotypes in young children with fragile X syndrome., Proceedings of the National Academy of Sciences of the United States of America 114 (2017) 10767–10772. 10.1073/pnas.1620994114.

[21] W. Cheng, Y. Zhang, M. Liu, Identification of Subtypes of HCC Using Bioinformatics and the Hepatocyte Differentiation Model., Methods in Molecular Biology (Clifton, N.J.) 2544 (2022) 253–258. 10.1007/978-1-0716-2557-6_18.

[22] T.S.C. Hinks, T. Brown, L.C.K. Lau, H. Rupani, C. Barber, S. Elliott, J.A. Ward, J. Ono, S. Ohta, K. Izuhara, R. Djukanović, R.J. Kurukulaaratchy, A. Chauhan, P.H. Howarth, Multidimensional endotyping in patients with severe asthma reveals inflammatory heterogeneity in matrix metalloproteinases and chitinase 3-like protein 1., The Journal of Allergy and Clinical Immunology 138 (2016) 61–75. 10.1016/j.jaci.2015.11.020.

[23] T.O.F. Ba-Dhfari, Hypothesis formulation in medical records space, PhD thesis, University of Manchester, 2017.

[24] M. Herrero-Zazo, T. Fitzgerald, V. Taylor, H. Street, A.N. Chaudhry, J.R. Bradley, E. Birney, V.L. Keevil, Using machine learning to model older adult inpatient trajectories from electronic health records data, iScience 26 (2023) 105876. 10.1016/j.isci.2022.105876.

[25] C.H. Waddington, The Strategy of Genes, Routledge, 2014.

[26] S. Wright, The roles of mutation, inbreeding, crossbreeding, and selection in evolution, Proceedings of the VI International Congress of Genetrics 1 (1932) 356–366.

[27] H.F. Nijhout, J. Best, M. Reed, Escape from homeostasis, Mathematical Biosciences 257 (2014) 104–110.

[28] American Cancer Society, Key Statistics for Chronic Lymphocytic Leukemia, (n.d.).

[29] C. Shustik, I. Bence-Bruckler, R. Delage, C.J. Owen, C.L. Toze, S. Coutre, Advances in the treatment of relapsed/refractory chronic lymphocytic leukemia, Annals of Hematology 96 (2017) 1185–1196. 10.1007/s00277-017-2982-1.

[30] N.E. Kay, P.J. Hampel, D.L. Van Dyke, S.A. Parikh, CLL update 2022: A continuing evolution in care, Blood Reviews 54 (2022) 100930. 10.1016/j.blre.2022.100930.

[31] H. Duzkale, C.D. Schweighofer, K.R. Coombes, L.L. Barron, A. Ferrajoli, S. O’Brien, W.G. Wierda, J. Pfeifer, T. Majewski, B.A. Czerniak, LDOC1 mRNA is differentially expressed in chronic lymphocytic leukemia and predicts overall survival in untreated patients, Blood 117 (2011) 4076–4084.

[32] L.V. Abruzzo, C.D. Herling, G.A. Calin, C. Oakes, L.L. Barron, H.E. Banks, V. Katju, M.J. Keating, K.R. Coombes, Trisomy 12 chronic lymphocytic leukemia expresses a unique set of activated and targetable pathways, Haematologica 103 (2018) 2069–2078. 10.3324/haematol.2018.190132.

[33] M.R. Zucker, L.V. Abruzzo, C.D. Herling, L.L. Barron, M.J. Keating, Z.B. Abrams, N. Heerema, K.R. Coombes, Inferring clonal heterogeneity in cancer using SNP arrays and whole genome sequencing, Bioinformatics 35 (2019) 2924–2931. 10.1093/bioinformatics/btz057.

[34] C.D. Herling, K.R. Coombes, A. Benner, J. Bloehdorn, L.L. Barron, Z.B. Abrams, T. Majewski, J.E. Bondaruk, J. Bahlo, K. Fischer, M. Hallek, S. Stilgenbauer, B.A. Czerniak, C.C. Oakes, A. Ferrajoli, M.J. Keating, L.V. Abruzzo, Time-to-progression after front-line fludarabine, cyclophosphamide, and rituximab chemoimmunotherapy for chronic lymphocytic leukaemia: A retrospective, multicohort study, Lancet Oncol 20 (2019) 1576–1586. 10.1016/S1470-2045(19)30503-0.

[35] R Core Team, R: A Language and Environment for Statistical Computing, R Foundation for Statistical Computing, Vienna, Austria, 2022.

[36] T.M. Therneau, P.M. Grambsch, Modeling Survival Data: Extending the Cox Model, Springer, New York, NY, 2000. 10.1007/978-1-4757-3294-8.

[37] P.J. Rousseeuw, L. Kaufman, Finding groups in data, Hoboken: Wiley Online Library (1990).

[38] P.J. Rousseeuw, Silhouettes: A graphical aid to the interpretation and validation of cluster analysis, Journal of Computational and Applied Mathematics 20 (1987) 53–65.

[39] Z.B. Abrams, C.E. Coombes, S. Li, K.R. Coombes, Mercator: A Pipeline For Multi-Method, Unsupervised Visualization And Distance Generation, Bioinformatics (2021). 10.1093/bioinformatics/btab037.

[40] S.-S. Choi, S.-H. Cha, C.C. Tappert, A survey of binary similarity and distance measures, Journal of Systemics, Cybernetics and Informatics 8 (2010) 43–48.

[41] B. Efron, R. Tibshirani, Empirical bayes methods and false discovery rates for microarrays, Genet Epidemiol 23 (2002) 70–86. 10.1002/gepi.1124.

[42] C.I. Amaya-Chanaga, L.Z. Rassenti, Biomarkers in chronic lymphocytic leukemia: Clinical applications and prognostic markers, Best Practice & Research. Clinical Haematology 29 (2016) 79–89. 10.1016/j.beha.2016.08.005.

[43] M. Crespo, F. Bosch, N. Villamor, B. Bellosillo, D. Colomer, M. Rozman, S. Marcé, A. López-Guillermo, E. Campo, E. Montserrat, ZAP-70 expression as a surrogate for immunoglobulin-variable-region mutations in chronic lymphocytic leukemia, The New England Journal of Medicine 348 (2003) 1764–1775. 10.1056/NEJMoa023143.

[44] R.N. Damle, T. Wasil, F. Fais, F. Ghiotto, A. Valetto, S.L. Allen, A. Buchbinder, D. Budman, K. Dittmar, J. Kolitz, S.M. Lichtman, P. Schulman, V.P. Vinciguerra, K.R. Rai, M. Ferrarini, N. Chiorazzi, Ig V gene mutation status and CD38 expression as novel prognostic indicators in chronic lymphocytic leukemia, Blood 94 (1999) 1840–7.

[45] D. Capello, A. Guarini, E. Berra, F.R. Mauro, D. Rossi, E. Ghia, M. Cerri, J. Logan, R. Foà, G. Gaidano, Evidence of biased immunoglobulin variable gene usage in highly stable B-cell chronic lymphocytic leukemia, Leukemia 18 (2004) 1941–1947. 10.1038/sj.leu.2403537.

[46] B. Simonsson, L. Wibell, K. Nilsson, Beta 2-microglobulin in chronic lymphocytic leukaemia, Scandinavian Journal of Haematology 24 (1980) 174–180. 10.1111/j.1600-0609.1980.tb02364.x.

[47] H. Döhner, S. Stilgenbauer, K. Döhner, M. Bentz, P. Lichter, Chromosome aberrations in B-cell chronic lymphocytic leukemia: Reassessment based on molecular cytogenetic analysis, Journal of Molecular Medicine 77 (1999) 266–281.

[48] E. Matutes, A. Polliack, Morphological and immunophenotypic features of chronic lymphocytic leukemia, Reviews in Clinical and Experimental Hematology 4 (2000) 22–47. 10.1046/j.1468-0734.2000.00002.x.

[49] A. Jenner, R. Aogo, V. Crowe, X. Deng, A. Smith, P. Morel, C. Davis, A. Smith, M. Craig, COVID-19 virtual patient cohort suggests immune mechanisms driving disease outcomes, PLoS Pathogens 17 (2021). 10.1371/journal.ppat.1009753.

