## Supplementary figures and images for "Topological Structures in the Space of Treatment-Naïve Patients With Chronic Lymphocytic Leukemia"

### Suppplemental Figure 1

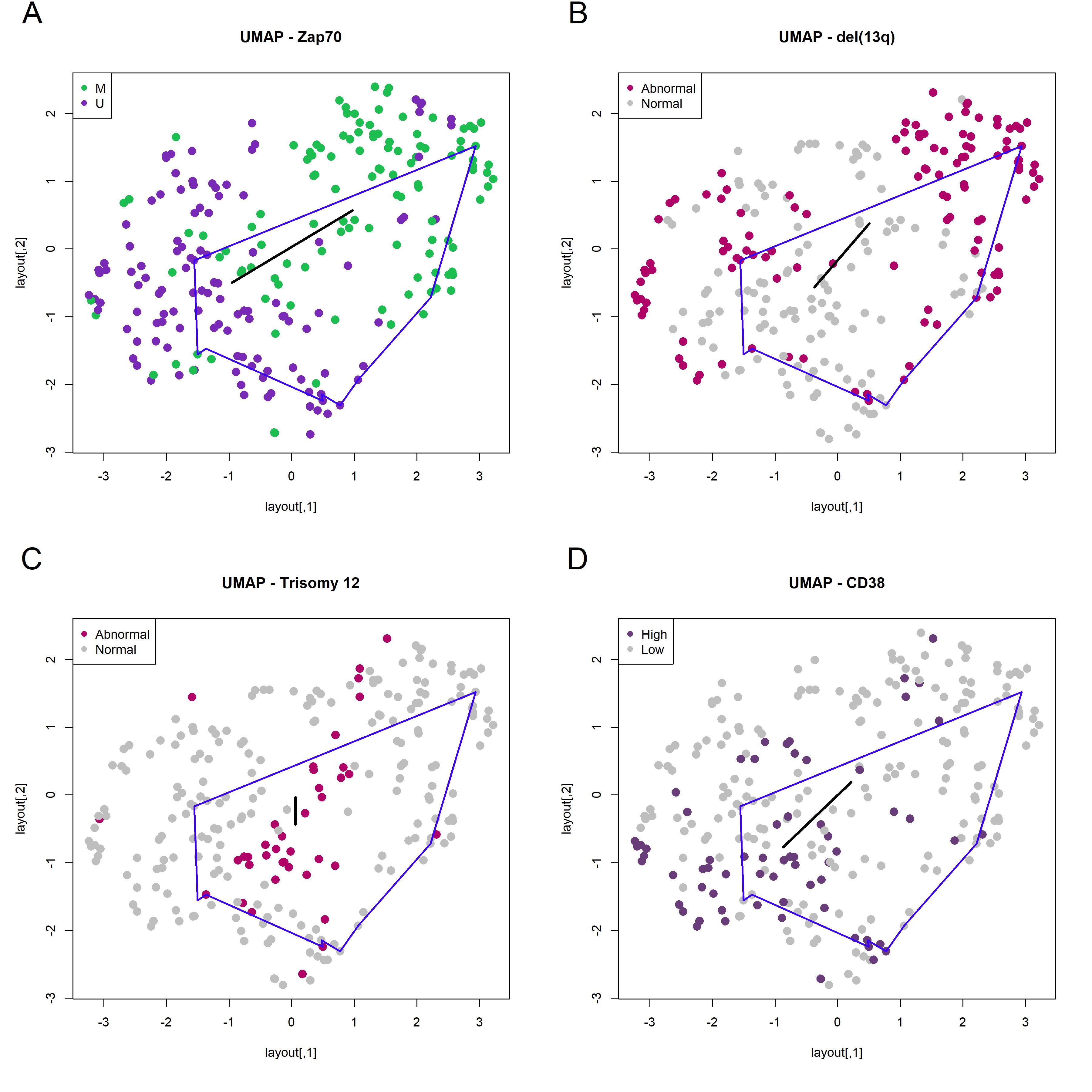

### Suppplemental Figure 2

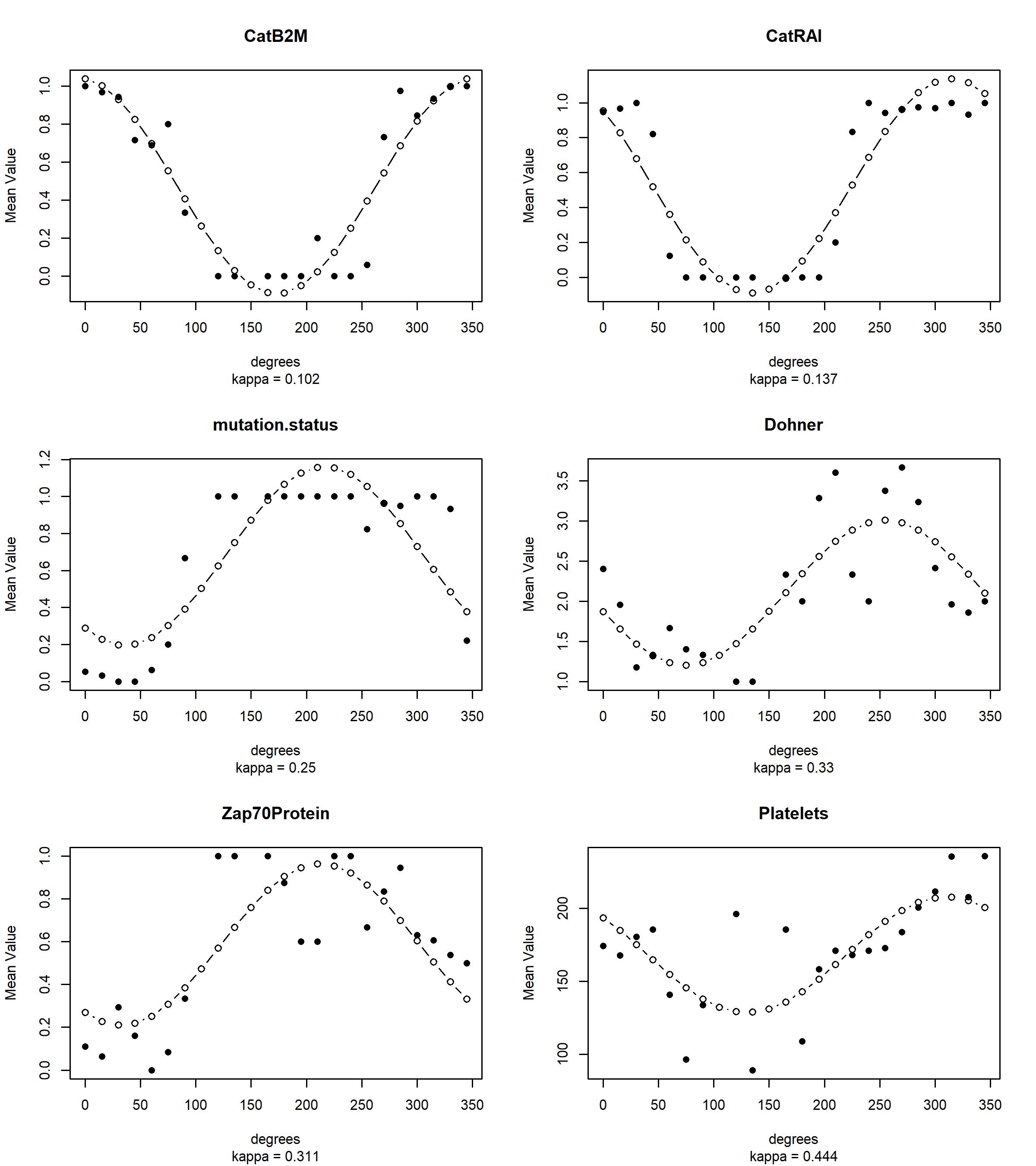

### Suppplemental Figure 3

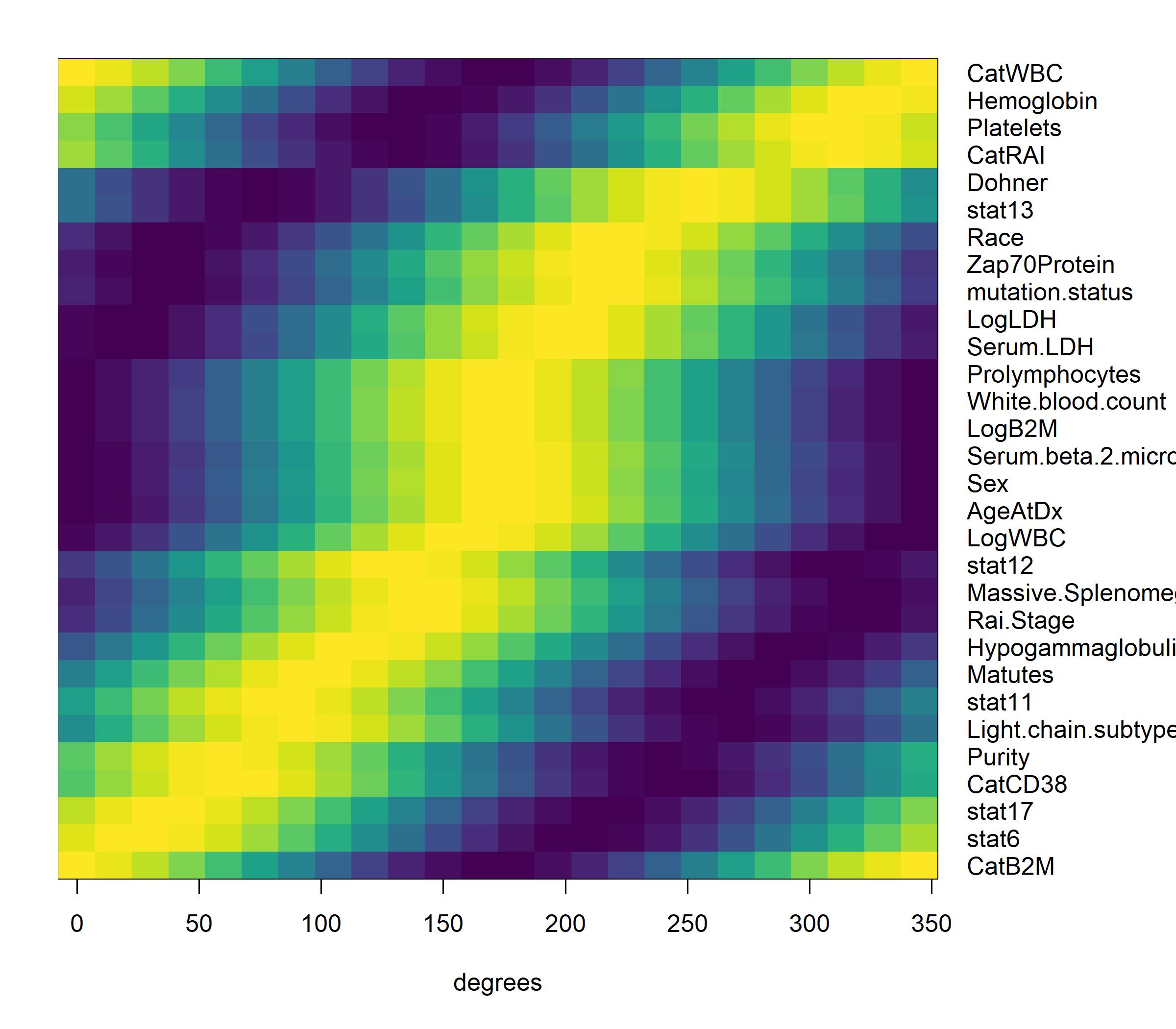

### Suppplemental Figure 4

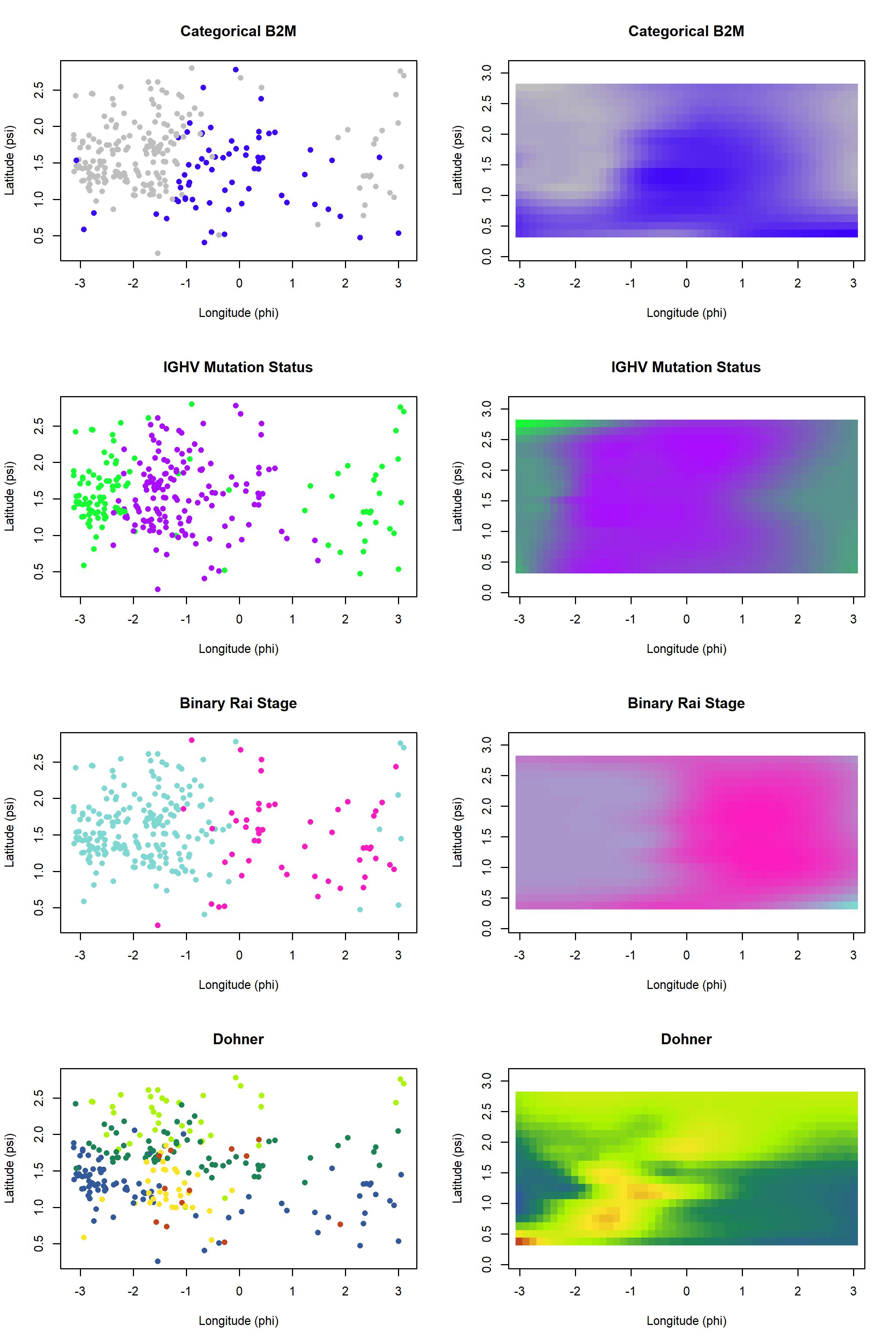

### Suppplemental Figure 5

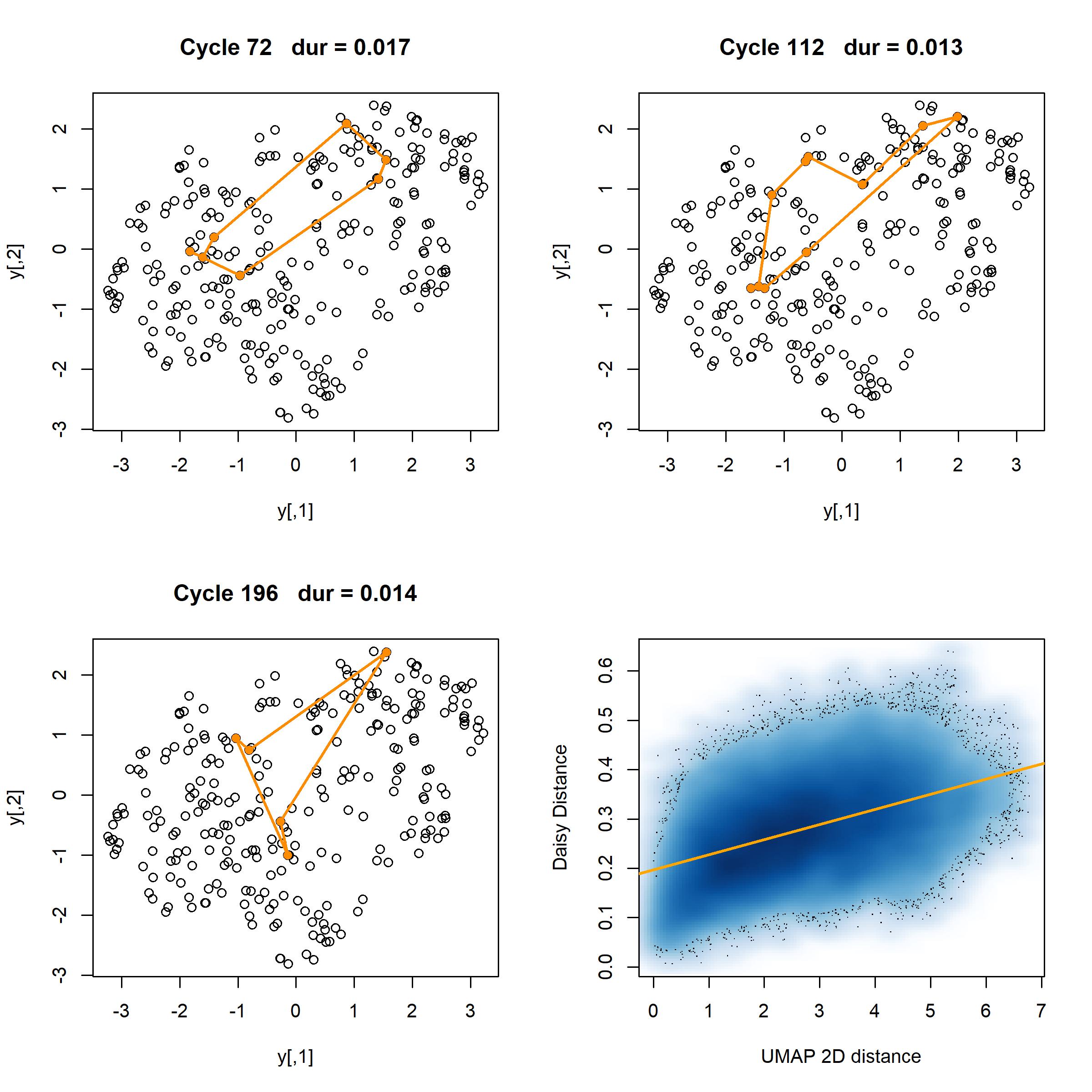
